# Direct synthesis of self-organized blastocyst-like cysts derived from human pluripotent stem cells

**DOI:** 10.1101/831487

**Authors:** Xiaopeng Wen, Shiho Terada, Koki Yoshimoto, Ken-ichiro Kamei

## Abstract

We introduce a simple, robust and scalable method to generate self-organized blastocyst-like cysts (soBLCs) from human pluripotent stem cells (hPSCs). We use a copolymer hydrogel of poly(*N*-isopropylacrylamide) and poly(ethylene glycol) (PNIPAAm-PEG). hPSC aggregates with a diameter of approximately 117.2 ± 5.1 µm are cultured in a medium supplemented with a hydrogel and a serum for three days. Molecular signatures in the medium revealed the generation of trophoblasts and inner cell mass at specific positions in the soBLCs.

## Main

A blastocyst is a cyst developed from the fertilized egg of a mammal approximately 5 to 8 days after fertilization, and it consists of a trophoblast and an inner cell mass (ICM) with a blastocyst cavity. This structure, with its unique cell positioning, is essential for further embryonic development for tissue-specific lineage cell differentiation, and subsequently organ and body structure formation. Therefore, a blastocyst will not only help study the early development of a cell but can also be used in drug discovery for pregnancy failure (e.g., repeated implantation failure^1^) and birth deficiency prevention^2^. However, especially in human beings, it is very difficult to obtain blastocysts from fertilized eggs for fundamental research as well as industrial usage due to limited cell sources / donors and ethical concerns.

Recent studies have reported methods to generate blastocyst-like structures for experimental mouse models^3,4^. However, these studies used and digested blastocysts from fertilized eggs to obtain embryonic stem cells (ESCs)^5^ and trophoblast stem cells (TSCs). They re-assembled these ESCs and TSCs to generate blastocyst-like cysts (BLCs). Thus, it is still challenging to generate BLCs directly from pluripotent stem cells alone (PSCs; such as ESCs and induced pluripotent stem cells [iPSCs]^6,7^). In a suspension cell culture, PSCs exhibit embryoid body (EB) formation with cells of three germ layers (e.g., endoderm, mesoderm and ectoderm) under differentiation conditions, or cell spheroids with self-renewing human PSC (hPSC). However, any such reported suspension cell culture did not produce blastocyst-like cysts.

A new method to generate BLCs from hESCs has been reported recently, but it requires special apparatus, such as a microfluidic device.^8^ This necessitates an alternative way to culture PSCs and produce BLCs in a simple, robust and scalable way that could be used in biology research laboratories and future drug discovery applications.

Here we develop a method to directly form self-organized BLCs (soBLCs) from hPSCs in a hydrogel medium through three-dimensional (3D) cell culture (**Figure 1a**). This method uses a thermo-responsive hydrogel [HG; a copolymer of poly(*N*-isopropylacrylamide) and poly(ethylene glycol) (PNIPAAm-PEG)]^9,10^, which holds hPSC aggregates with mild physical stimuli, without adhering with the cells. Additionally, because it can perform a sol–gel transition via temperature, the HG allows the mixing and harvesting of cell aggregates from the solution at a low temperature (< 20 °C). However, cell culture in a gel medium at high temperatures (37 °C). In comparison, we tested collagen and agarose hydrogels for BLC generation (**Figure S1**). For the differentiation of a part of the hPSC aggregates to the trophoblast lineage, the HG was dissolved in DMEM supplemented with 10%(v/v) fetal bovine serum (FBS). For comparison, a TeSR-E8 hPSC self-renewal medium^11^ was also mixed with 10%(w/v) HG. To increase the reproducibility, single-cell-dissociated hPSCs were transferred to AggreWell^12,13^ for cell aggregates with controlled uniform sizes. One day after the formation, the hPSC aggregates were mixed in the HG and cultured for 3–5 d. To visualize the expression of octamer-binding transcription factor 4 (OCT4; or POU domain, class 5, transcription factor 1 [POU5F1]) ICM marker, *OCT4* promoter-driven KhES1 hESCs with an enhanced green fluorescent protein (EGFP) were used (named K1-OCT4-EGFP).^14^

**Figure 1.**
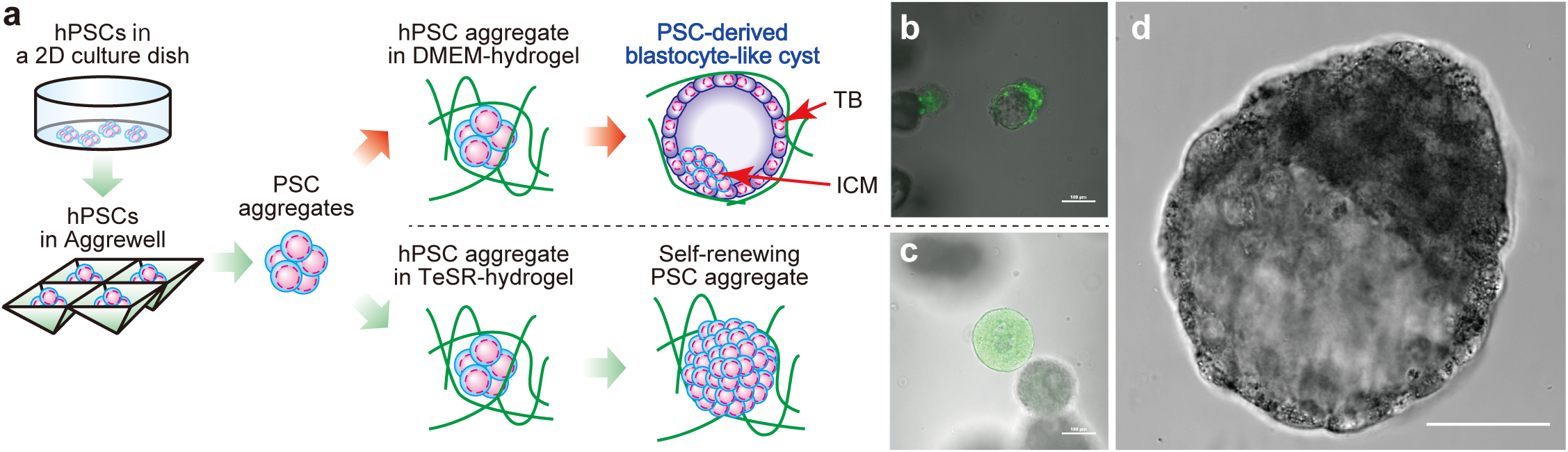
Principle of generating self-organized blastocyst-like cell cysts (soBLCs) derived from human pluripotent stem cells (hPSCs). **a**, Schematic of experimental process of generating blastocyst-like cysts from hPSCs. Hydrogel with self-renewing hPSC culture medium (e.g., TeSR-E8) provides spheroid cells. Hydrogel dissolved in DMEM with 10%(v/v) FBS provides cell aggregates with a small cavity inside, which resembles a blastocyst. TB; trophoblasts. ICM; inner cell mass. **b**, An overlaying micrograph of DIC and green fluorescence of a self-organized blastocyst-like cyst (soBLC) derived from K1-OCT4- EGFP cells in a hydrogel containing DMEM with 10%(v/v) FBS. Scale bar represents 100 µm. **c**, An overlaying micrograph of DIC and green fluorescence of an aggregate of K1- OCT4-EGFP cells in hydrogel contain TeSR-E8 medium. These cell aggregates did not have cavities inside. Scale bar represents 100 µm. **d**, A typical DIC micrograph of soBLC. Scale bar represents 50 µm.

In a first, 38.8% of hPSC aggregates (21 out of 54 aggregates) cultured in the hydrogel mixed with DMEM/FBS conditions formed cell aggregates with a cavity, which has a structure closely resembling a blastocyst. On the one hand, a small portion of the cell aggregates expressed the *OCT4* promoter-driven EGFP. The outer cell layer, however, did not express EGFP adequately (**Figure 1b**). On the other hand, K1-OCT4-EGFP cells cultured in TeSR-E8 medium-mixed hydrogels showed cell spheroids that strongly expressed EGFP— meaning that cultured hPSCs maintain its stemness in the tested hydrogels (**Figure 1c**). In fact, these spheroids contain small cavities; however, most of the cells retained the EGFP expression. This result indicates that the self-renewing stimulants in the TeSR-E8 medium strongly retained the stemness of the cultured hPSCs, even in hydrogel. This result is in agreement with that of our previous report^9^. These results indicate that hydrogel conditions along with the serum facilitate the formation of cell aggregates with blastocyst-like structure (**Figure 1d**).

We then evaluated the molecular signatures as indication of a blastocyst (**Figure 2**). To this end, first, we observed the expression of caudal homeobox 2 (*CDX2*^15^), which trophoblasts of a blastocyst specifically expressed (**Figure 2a**). We also evaluated the expression of *OCT4*. While the self-renewing K1-OCT4-EGFP cells expressed OCT4 but not the *CDX2* gene, the obtained hBLCs increased the expression of *CDX2* (6.2 folds) and reduced that of *OCT4* (0.44 folds). We also observed other markers for the trophoblasts [e.g., Leukemia Inhibitory Factor Receptor (*LIFR*), Keratin 8 (*KRT8*), and GATA binding protein 3 (*GATA3*)] confirmed that those markers were specifically found in soBLCs (2.27, 8.65, 11.1 folds compared with self-renewing K1 hESCs, respectively), and the marker of ICM (e.g., *NANOG*) were reduced in soBLCs (0.21 folds). These results also imply that the blastocyst cells were differentiated by hPSCs in the cysts. To confirm the cellular distribution of trophoblasts and ICM cells in soBLCs, we observed the expression patterns of OCT4 and CDX2 in soBLCs via whole-mount immunocytochemistry (**Figure 2b**). On the one hand, the cells located in the outer layer expressed CDX2, but those inside the cysts did not. On the other hand, the cells both inside and outside of the cyst expressed OCT4. These results indicate formating a blastocyst-like structure. To evaluate the cellular population within the soBLCs, we conducted flow cytometry to gauge the expressions of SSEA-1 and SSEA-4, which are also markers of trophoblast and ICM cells, respectively^16–18^ (**Figure 2c**). In the soBLCs, the percentages of SSEA-1^+^: SSEA-4^-^ was 0.593%, while the self-renewing hPSC spheroids contain only the 0.055% of SSEA-1^+^: SSEA-4^-^ cells.

**Figure 2.**
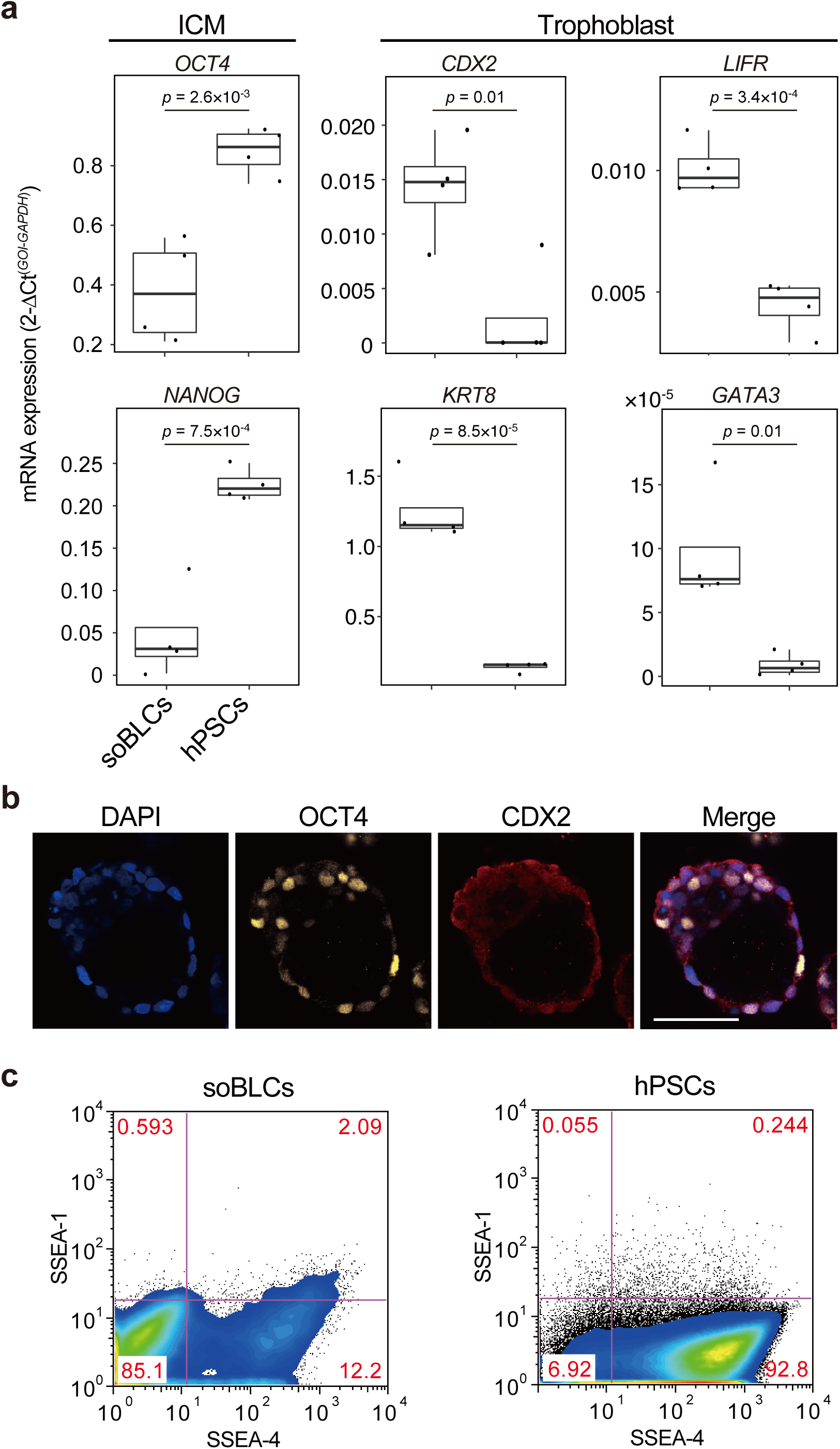
hPSC-derived BLCs with identical structures and molecular signature of a blastocyst. **a**, Box plots of gene expressions associated with inner cell mass (ICM; *OCT4* and *NANOG*) and trophoblasts (*CDX2, LIFR, KRT8*, and *GATA3*). Center lines of boxplots are the medians; box limits indicate the 25th and 75th percentiles; whiskers indicate the 9th and 91th percentiles. *P* values were estimated by two-tailed Student’s t-test. **b**, Micrographs of immunocytochemistry to observe the expressions of OCT4 and CDX2. DAPI was used for nuclear staining. Scale bar is 100 µm. **c**, Flow cytometric analysis to evaluate the cellular populations of soBLCs and self-renewing hPSCs according to the expressions of SSEA-1 and SSEA-4. The typical graphs of flow cytometry from the experiments, which were repeated at least thrice, are shown here.

In conclusion, we established a simple and robust method to generate human soBLCs from only hPSCs. The hydrogel enables better environments for the formation of soBLCs, which suggests that a certain physical property of the gel plays an important role in the soBLC formation and not the fluid property. Recently, some methods for BLC production have been. However, they need to use *in vitro* fertilized mice eggs and dissociate them. We envision that this method could be improved further to develop clinically relevant and chemically defined conditions, with the goal of understanding the basis of early embryonic development. By using PSCs of other experimental animals (e.g. mouse and rat), we should be able to evaluate the applicability of our methods for BLC generation. This method could be extended to other species to understand the fundamentals of early embryonic development, by comparing multiple species without using their embryos, and to high-throughput screening in drug discovery for pregnancy failure prevention, as well as birth defects.

## Supporting information

Supplemental informations

## Acknowledgements

We thank Dr. Kouichi Hasegawa and Dr. Dan Ohtan Wang for critical discussion. Funding was generously provided by the Japan Society for the Promotion of Science (JSPS; 17H02083). The WPI-iCeMS is supported by the World Premier International Research Centre Initiative (WPI), MEXT, Japan.

## Author contributions

X.W. and K.K. conceptualized the work. X.W., K.Y. and S.T. performed the experiments. All the authors contributed to data analysis, discussion, and interpretation. X.W. and K.K. wrote and revised the manuscript with input from all authors.

## Methods

### Self-organization of blastocyst-like cyst (soBLC) from hPSCs

hESCs were used according to the guidelines of the ethical committee of Kyoto University. K1-OCT4-EGFP was obtained from Eihachiro Kawase.^14^ Cultured on Matrigel-coated dishes in TeSR-E8 medium, the hPSCs were trypsinized and collected in a 15-mL tube. The dissociated hPSCs were resuspended in the Dulbecco’s modified Eagle medium (DMEM) supplemented with 10% (v/v) FBS (Cell Culture Bioscience), 1% (v/v) non-essential amino acids (NEAA; Thermo Fisher Scientific), and 1% (v/v) penicillin/streptomycin (PS; Thermo Fisher Scientific), then transferred into Aggrewell 400 (Stem Cell Technologies) at 6 × 10^5^ cells per well. After culturing in an incubator for 24 h at 37 °C with 5% (v/v) CO_2_, the hPSC aggregates were resuspended in blastocyst formation medium {DMEM supplemented with 10% (v/v) FBS, 1% (v/v) NEAA, and 1% (v/v) PS, 10%[w/v] HG (Mebiol Inc., Hiratsuka, Japan)} or TeSR- E8/HG medium (TeSR-E8 medium supplemented with 10%[w/v] HG and 10 µM Y-27632 [Wako Chemicals]) at 4 °C. 500 µL of an hPSC aggregate suspension with HG was transferred into a well of a 6-well plate, and then the appropriate culture medium was added at 37 °C. The medium was changed daily, and the cells were maintained in an incubator at 37 °C with 5% (v/v) CO_2_.

