## Supplemental informations for "Direct synthesis of self-organized blastocyst-like cysts derived from human pluripotent stem cells"

**Methods**

**hPSC culture.** hESCs were used according to the guidelines provided by the ethical committee of Kyoto University. K1-OCT4-EGFP was obtained from Eihachiro Kawase.^1^ Prior to culturing, hESC-certified Matrigel (Corning, NY, USA) was diluted with a DMEM/F12 medium (Sigma-Aldrich, St. Louis, MO, USA) at 1.3% (v/v) and was coated on a culture dish. Matrigel was incubated in a dish for 24 h at 4°C. Then, excess Matrigel was removed, and the coated dish was washed with a fresh DMEM/F12 medium. A TeSR-E8 medium (Stem Cell Technologies, Vancouver, Canada) was used for the daily culturing of hPSCs. For passaging, cells were dissociated with TrypLE Express (Life Technologies, Carlsbad, CA, USA) for 3 min at 37 °C, and then harvested. A cell strainer was used to remove undesired cell aggregates from the cell suspension. The cells were centrifuged at 200 × *g* for 3 min and resuspended in the TeSR-E8 medium. The cells were counted using a NucleoCounter NC-200 (Chemetec, Baton Rouge, LA, USA). A TeSR-E8 medium containing 10 µM of Y27632 ROCK inhibitor (Wako, Osaka, Japan) was used to prevent the apoptosis of the dissociated hPSCs, on day 1. A TeSR-E8 medium without Y27632 was used on the subsequent days, with daily medium change. The cells were maintained in an incubator at 37°C with 5% (v/v) CO_2_.

**Self-organization of blastocyst-like cyst (soBLC) from hPSCs.** hPSCs cultured on Matrigel-coated dishes with a TeSR-E8 medium were trypsinized and collected in a 15-mL tube. The dissociated hPSCs were resuspended in DMEM supplemented with 10% (v/v) fetal bovine serum (FBS; Cell Culture Bioscience), 1% (v/v) non-essential amino acids (Thermo Fisher Scientific), and 1% (v/v) penicillin/streptomycin (Thermo Fisher Scientific), and then transferred to Aggrewell 400 (Stem Cell Technologies) at 6 × 10^5^ cells per well. After culturing in an incubator for 24 h at 37°C with 5% (v/v) CO_2_, the hPSC aggregates were resuspended in the blastocyst formation medium {DMEM supplemented with 10% (v/v) fetal bovine serum (FBS; Cell Culture Bioscience), 1% (v/v) non-essential amino acids (Thermo Fisher Scientific), and 1% (v/v) penicillin/streptomycin (Thermo Fisher Scientific), 10%[w/v] PNIPAAm-β-PEG hydrogel (HG; Mebiol Inc., Hiratsuka, Japan)} or a TeSR-E8/HG medium (TeSR-E8 medium supplemented with 10%[w/v] HG and 10 µM Y-27632) at 4°C. 500 µL of an hPSC aggregate suspension with HG was transferred into the well of a 6-well plate, and then the corresponding culture medium was added at 37°C. The medium was changed on a daily basis, and the cells were maintained at 37°C with 5% (v/v) CO_2_ in an incubator.

**Cell aggregate culture in collagen hydrogel.** Cell aggregate culture in collagen or agarose was performed following the section of “Self-organization of blastocyst-like cyst (soBLC) from hPSCs,” except the use of collagen (Nippi) and agarose as alternatives to the HG. 10%(w/v) of collagen and 2%(w/v) of agarose were used for the cell aggregate culture.

**Quantitative RT-PCR.** Total RNA was purified using an RNeasy Mini Kit (Qiagen, Hilden, Germany). For the positive control of human normal liver, Human Total Liver RNA was purchased from TaKaRa Bio. Total RNA (2.5 µg) was reverse-transcribed to generate cDNA using PrimeScript RT master mix. A reaction mixture (21 µL) of 20 ng cDNA, 12.5 µL SYBR Premix Ex Taq II (Tli RNaseH Plus; TaKaRa Bio), and 0.5 µL ROX reference dye was introduced into a tube. The PCR conditions included initial incubation at 95°C for 30 s, followed by 40 cycles of 95°C for 5 s and then 60°C for 31 s on an Applied Biosystems 7300 real-time PCR system.

**Immunocytochemistry.** Cells were fixed with 4%(v/v) paraformaldehyde in PBS for 20 min at 25°C, and then permeabilized with 0.1%(w/v) sodium dodecyl sulfate in PBS for 16 h at 25°C. Subsequently, the cells were blocked in PBS (5%[v/v] normal goat serum, 5%[v/v] normal donkey serum, 3%[w/v] bovine serum albumin, 0.1%[v/v] Tween-20) at 4°C for 16 h, and then incubated at 4°C for 16 h, with the primary antibody (anti-human CDX2 antibody ab76541; Abcam, Cambridge, UK; anti-human OCT3/4 antibody C-10; Santa Cruz Biotechnology, CA, USA) in PBS with 0.5% Triton X-100. The cells were then incubated at 37°C for 60 min with a secondary antibody (AlexaFluor 488 Donkey anti-rabbit IgG, 1:1000; Jackson ImmunoResearch, West Grove, PA, USA) in a blocking buffer prior to the final incubation with 300 nM of 4′,6-diamidino-2-phenylindole (DAPI) at 25°C for 30 min.

**Image acquisition**. The sample containing the cells was placed on the stage of a Nikon A1Plus inverted confocal laser scanning fluorescence microscope equipped with a CFI Plan Apo VC 20× DIC objective lens (Nikon, Tokyo, Japan).

**Flow cytometry.** Cell aggregates were harvested with trypsin (0.25%) and rinsed twice with PBS prior to cell counting. For staining with antibodies, the cells were diluted to a final concentration of 1 × 10^7^ cells mL^−1^ in PBS supplemented with 2% fetal calf serum (FCS). Fluorescence-labeled antibodies (SSEA-1 and SSEA-4) were added and incubated at 25°C for 30 min. As a negative control, specific isotype controls were used. After removing the excess antibodies via centrifugation at 300 × *g* for 5 min, the cells were washed with PBS containing 2% FCS. The cell suspensions were applied to a FACS Canto II (BD Biosciences, Franklin Lakes, NJ, USA) for flow cytometric analysis. Data analysis was performed using FlowJo software (v9; FlowJo, LLC, Ashland, OR, USA).

**Supplementary Figures**

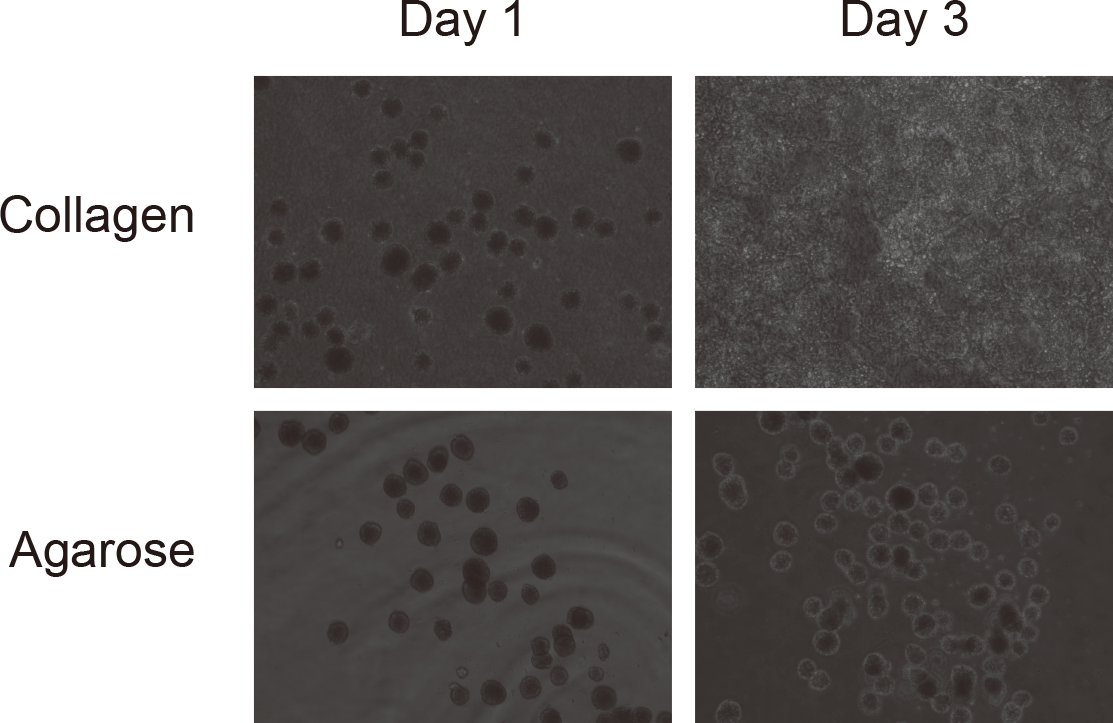

**Fig. S1 | Evaluation three-dimensional culture pf K1-OCT4-EGFP hESCs using collagen and agarose hydrogel.** K1-OCT4-EGFP hESCs formed spherical cell aggregates in both collagen and agarose hydrogel at Day 1. However, none of them could not form BLCs. No cell aggregates in agarose gel were observed. Cells in collagen hydrogel migrated into the gel and no longer formed spheroid cell aggregates at Day 3.
